# Cellular metabolic activity marker via selective turn-ON detection of serum albumin using an NBD-based fluoroprobe

**DOI:** 10.1101/732610

**Authors:** Tanoy Dutta, Kaushik Pal, Apurba Lal Koner

**Affiliations:** Department of Chemistry, Indian Institute of Science Education and Research Bhopal, Bhopal Bypass Road, Bhauri, Bhopal-462066, Madhya Pradesh, INDIA

**Keywords:** NBD dyes, Solvatochromism, Serum-binding, Live-cell imaging, Cancer-metabolism

## Abstract

A nitrobenzoxadiazole-based fluoroprobe (NBD-Bu) is designed to probe cellular metabolic activity in cancer and normal cells. NBD-Bu shows a significant fluorescence enhancement upon selective binding to serum albumin. The site specificity of NBD-Bu has been explored through a competitive displacement assay in the presence of site-specific markers such as warfarin and ibuprofen. Subsequently, high-resolution fluorescence microscopy results consolidated the potential of NBD-Bu for cellular imaging and detection of abnormal cellular metabolic activity.

## Introduction

Serum albumin (SA) is the most abundant water-soluble protein found in blood plasma occupying almost 60% of total blood plasma proteins.^1–2^ Human serum albumin (HSA) has 585 amino acid residues in a single polypeptide chain while bovine serum albumin (BSA), which is 76% homologous to HSA has 583 amino acid residues.^1^ It functions as a versatile transport protein, transporting small drug molecules, bile-salt, hormones, vitamins, metals and plays a significant role by contributing 80% to the maintenance of oncotic pressure between blood vessels and tissues.^3–4^ Moreover, albumins are primarily responsible for maintaining the pH of blood.^5–6^ SA is also used as an additive in the cell culture media as it enhances growth and cell viability.^7^ A high level of SA is triggered by severe dehydration and high protein diet while deficiency of the same causes dysfunction of the circulatory system, which inclines the detection of SA to be extremely important.^6^

Among the vastly employed and reported techniques in this purpose, cyclic voltammetry,^8–9^ circular dichroism spectroscopy,^10^ nuclear magnetic resonance spectroscopy,^11–12^ high-performance liquid chromatography, absorption spectroscopy, and mass spectrometry are quite useful. However, these methods are not as popular as fluorescence spectroscopy is because of their sophistication, poor sensitivity. Fluorescence spectroscopy is one of the widely used and efficient tools to study the interaction between drug molecules and SA because of its rapidness, good selectivity, as well as high sensitivity and especially real-time visualization by naked-eye.^13–14^ Several fluorescent dyes are already known for SA binding, but the binding site and stoichiometry is not explored in most of the cases.^14–17^ Quite a few numbers of fluorescent probes have been developed based on their sensitivity towards local polarity and viscosity, which causes a significant change in emissive states upon binding to the multiple hydrophobic pockets present in SA.^18^ Nevertheless, limitations arise due to the absorbance of the probes in the near-UV region resulting to the interference by the auto-fluorescence of protein molecules, change in the secondary structure of protein upon ligand binding and lower selectivity of the sensor while other relevant analytes are present.^19^ Nitrobenzoxadiazole (NBD) dyes are well explored in the literature for their interesting fluorogenic intramolecular charge-transfer (ICT) properties and utilized for sensing, and protein binding studies.^20^ NBD-labeled lipids are well-explored fluorescent probes for understanding membrane structure and dynamics, and have been widely used in both model systems and living cells.^21–22^

Herein, we developed a turn-ON electroneutral solvatofluorochromic probe, with excellent selectivity, specificity, and sensitivity towards microenvironment polarity and instantaneous binding efficiency towards BSA. The selectivity of NBD-Bu for BSA was assessed using several important proteins with varied structures and functions. Moreover, we have employed two site marker drugs *viz*. warfarin and ibuprofen to study the binding phenomena with our probe. Among the several hydrophobic binding cavities inside BSA, warfarin binds to site I while ibuprofen binds to site II.^23^ NBD-Bu shows a notable blue-shift in emission maxima upon binding to BSA and the binding constant is found to be higher than that of ibuprofen. The probe also survived the selectivity test even in the presence of other biologically relevant proteins. Furthermore, we were able to study the difference in metabolic activity of cancer cells as compared to normal cells using live-cell imaging.

## Result and Discussions

The NBD moiety is involved with the easy synthetic procedure and it is well-known for its sensitivity towards environment polarity thereby being a potential sensor for micellar as well as proteinaceous microenvironments. Based on previous reports, a short butyl chain was incorporated in the NBD moiety to impart better fluorescence property and appropriate lipophilicity for improved binding affinity with the hydrophobic pocket of the protein.^24^ Following the synthesis of NBD-Bu, it was characterized by NMR spectroscopy (Fig. S1-2) and mass spectrometry (Fig. S3). The optical purity was also confirmed by comparing the absorption and excitation spectra (Fig. S4). NBD-Bu contains an electron-rich nitrogen atom, tethered to a butyl chain and an electron deficient NO_2_ group. These two moieties are linked *via* an aromatic ring, which renders it to be a classical ICT dye. In general, ICT dyes are well responsive to the solvent polarity thereby exhibiting polarity-dependent photo-physical behavior.^25^

To validate the ICT properties of NBD-Bu (Scheme 1), we have performed a thorough spectroscopic investigation using different solvents with varying polarity. In both ground and excited state, the spectra shifted towards longer wavelength along with spectral broadening upon increasing solvent polarity (Fig. 1a-b, Table S1). Subsequently, the fluorescence lifetime of NBD-Bu (Fig. 1c) was found to be longer in the non-polar solvents compared to polar ones, which are mainly due to the presence of hydrogen bonding interaction between dye and solvent molecules originating from higher non-radiative decay rates in polar solvents. The emission maxima and fluorescence lifetime varied linearly (Fig. 1d, S5) with polarity consolidating the fact that NBD-Bu can be used for local environment sensing. Such a feature makes it an excellent candidate to sense the local micro-polarity inside the hydrophobic pockets of protein molecules.

**Scheme 1.**
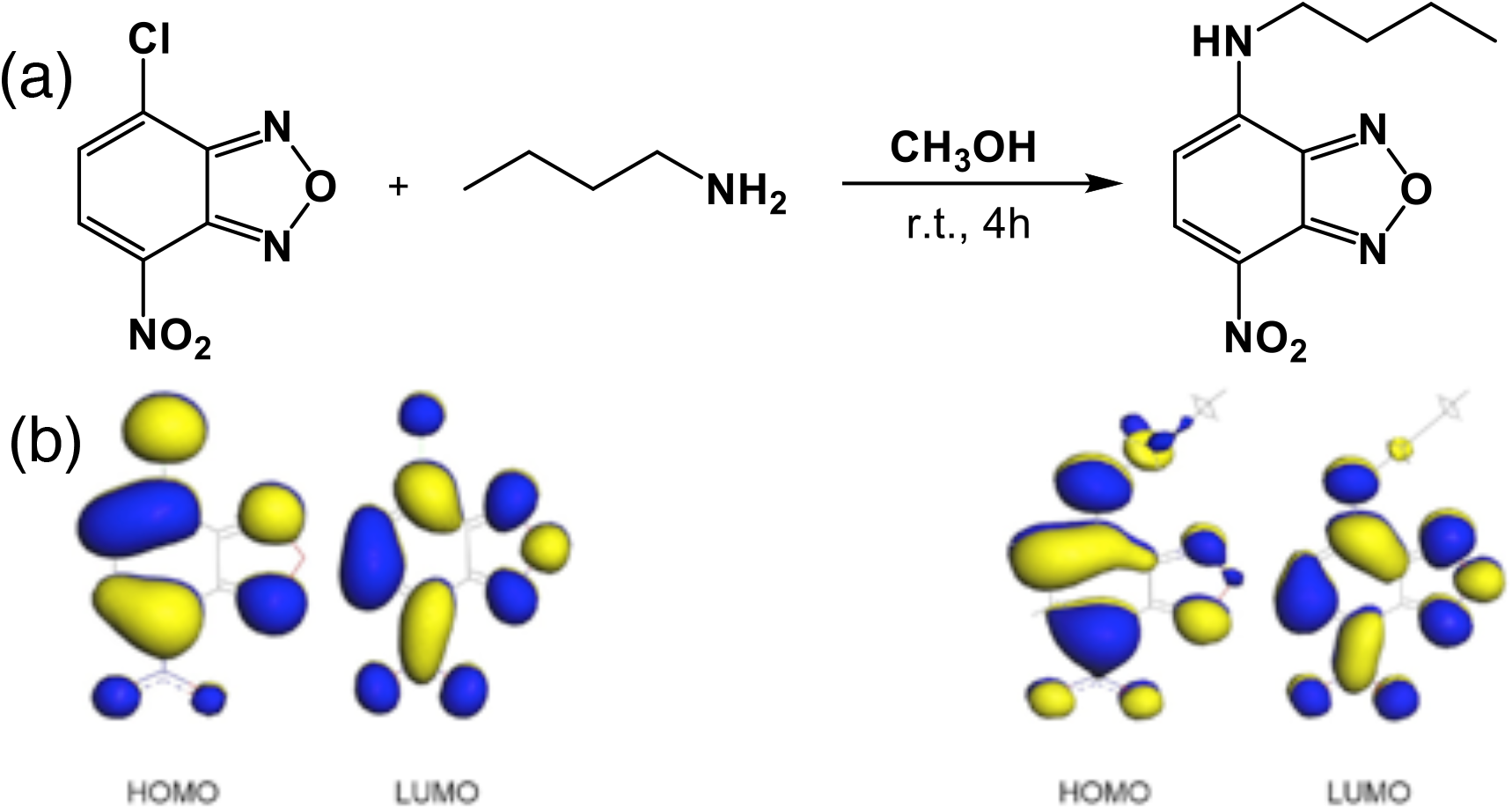
(a) Synthetic scheme of the fluoro-probe N-butyl-7-nitrobenzo[c][1,2,5]oxadiazol-4-amine (NBD-Bu), (b) frontier molecular orbital picture of NBD moiety before and after substitution

**Fig. 1.**
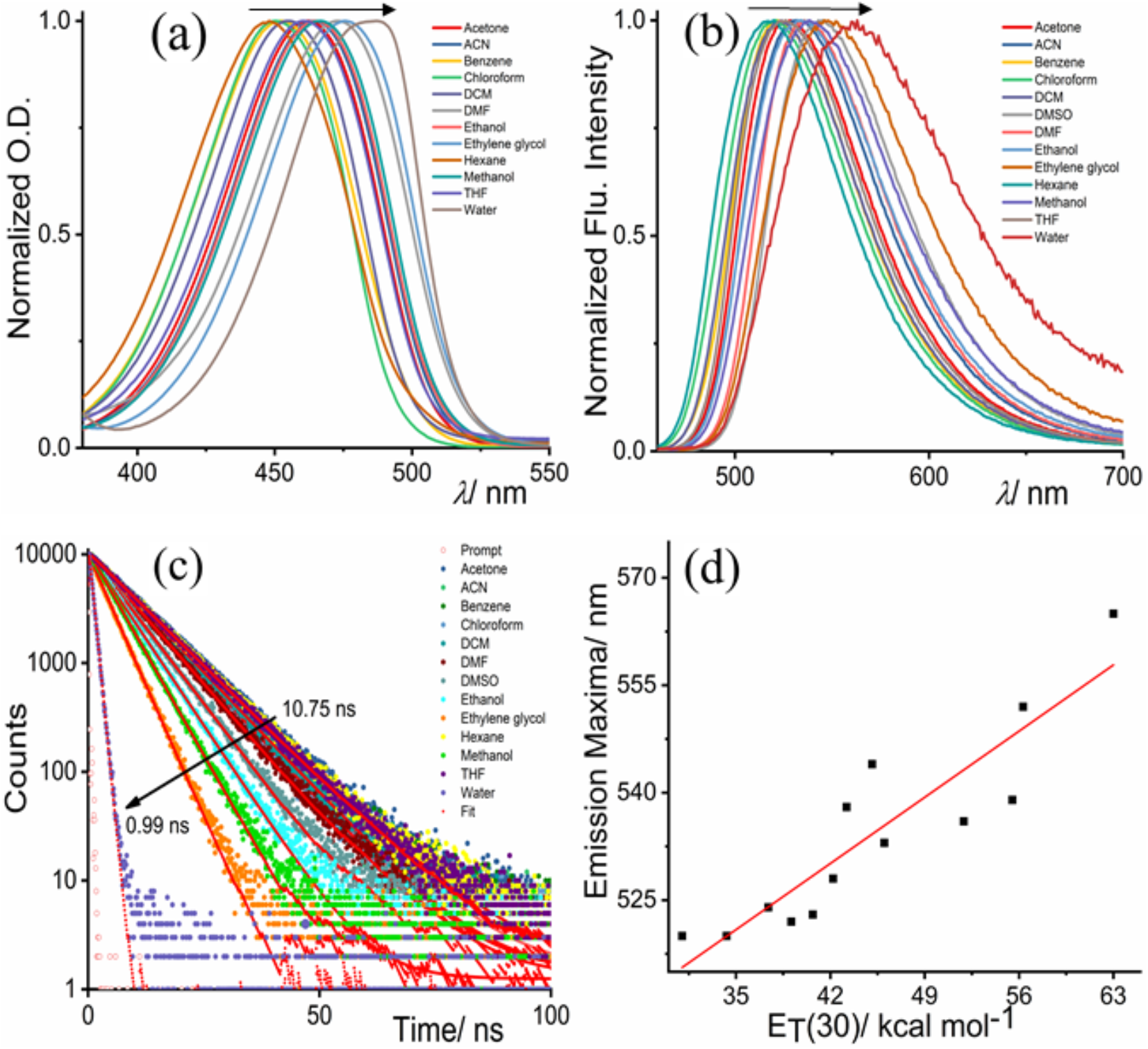
Solvent-dependent photophysical properties of NBD-Bu: (a) normalized UV-Vis. spectra with the varying solvent polarity, (b) normalized fluorescence spectra, (c) time-resolved fluorescence decay profile and (d) emission maxima vary linearly (*R*^2^ =0.82) with solvent polarity parameter. Arrows indicate the direction of low to high solvent polarity.

Considering the excellent solvatochromic property of NBD-Bu, we studied its absorption and fluorescence response for assessing hydrophobic pockets of proteins in PBS. NBD-Bu possesses good water solubility as well as excellent pH stability in the range of pH 2-9 (Fig. S8-9). The probe itself is weakly emissive in the aqueous medium; however, its fluorescence intensity increased gradually upon increasing BSA along with a significant 30 nm blue-shift without affecting the absorption maxima (Fig. 2a, S10 and Table S1). The emission maxima of NBD-Bu bound BSA was similar to the neat acetone which supports the previous report from Kudo *et al.*^26^ Upon addition of BSA, the fluorescence intensity increased thrice, instantaneously, triggering the propensity of detecting BSA in an expeditious manner.

**Fig. 2.**
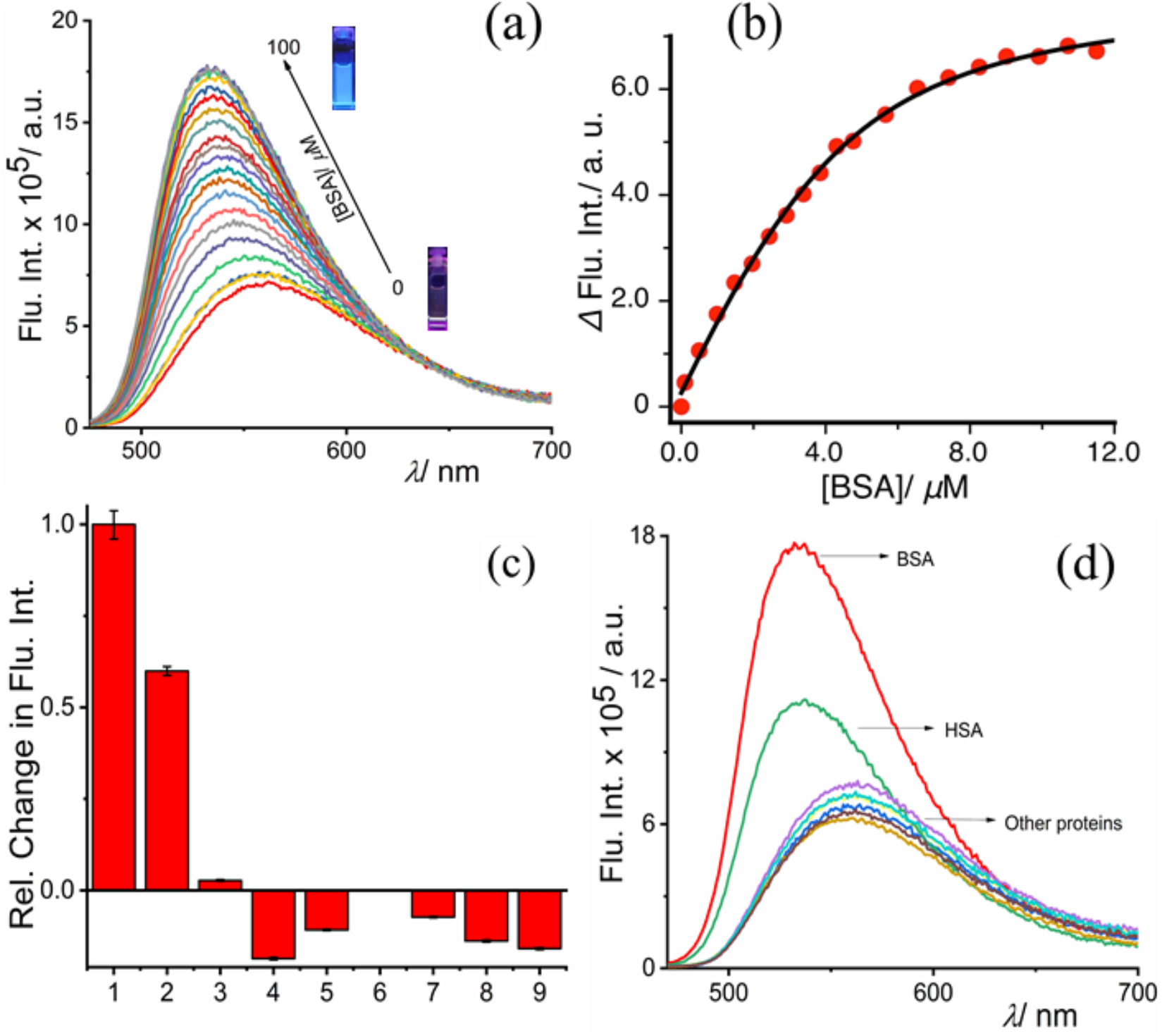
(a) Fluorescence titration of NBD-Bu (10 *µ*M) with increasing BSA (up to 100 µM) concentration (b) plot of fluorescence intensity of NBD-Bu with BSA concentration fitted with 1:1 binding equation, (c) relative change in the fluorescence intensities of NBD-Bu (10 *µ*M) in presence of other biologically relevant proteins (100 *µ*M): 1. BSA, 2. HSA, 3. Trypsin, 4. Lysozyme, 5. *α*-amylase, 6. *α*-Chymotrypsin, 7. RNAse, 8. DNAse and 9. *α*-lactalbumin, (d) the global analysis of fluorescent spectra of NBD-Bu with all proteins in PBS (pH 7.4).

This interesting result inspired us to investigate further the binding phenomena of NBD-Bu with BSA. As small molecules with similar van der Waals volume are known to bind with SA in 1:1 mode,^20^ hence, in this case, the binding curve was fitted (Fig. 2b) using the with 1:1 binding equation. The results showed a remarkably high binding constant, K= 8.3×10^5^ M^−1^. The limit of detection (LOD) was calculated to be 0.22 *µ*M (Fig. S11). The binding of NBD-Bu was further verified with fluorescence lifetime and time-resolved anisotropy analysis (Fig. S12-14). Furthermore, the selectivity, another important aspect which defines the applicability of a sensing probe, was assessed. In this regard, our probe was found to be non-responsive towards other physiologically relevant proteins with structural and functional diversity (Fig. 2c-d).

The slight interference in the presence of HSA can be attributed to its structural similarity with BSA. Hence, the selective behavior of the probe can be well acclaimed. Following up the binding studies of the probe with BSA, the site specificity within the hydrophobic pocket of the protein molecule was investigated. Ibuprofen and warfarin are well-explored drug molecules known to bind specifically to site-II and site-I, respectively. The competitive binding experiments (Fig. 3a-b) suggest that ibuprofen could replace the bound dye only to a certain extent indicating that NBD-Bu binds to the same site as that of ibuprofen and has a stronger binding affinity, too. Even 1 mM ibuprofen could only reduce the intensity up to ~25% along with a 10 nm red-shift. Interestingly, warfarin showed no effect upon similar treatment and the fluorescence intensity remained unchanged. This proves that NDB-Bu specifically binds to site-II of BSA without affecting its secondary structure (Fig. S15).

**Fig. 3.**
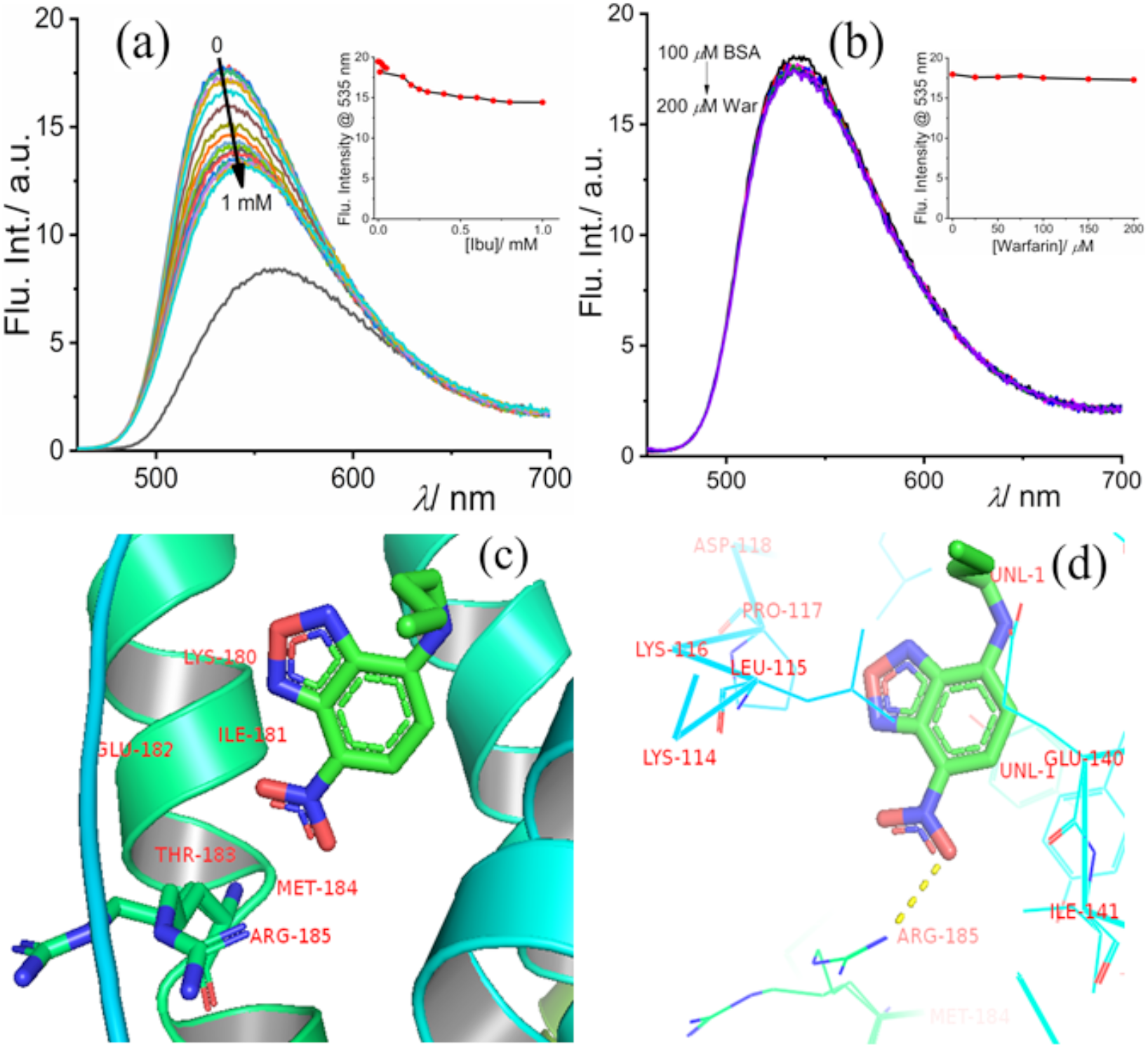
Competitive assay of NBD-Bu (10 *µ*M), pre-incubated with 100 *µ*M BSA, with increasing concentration of (a) ibuprofen and (b) warfarin in PBS; (inset: the intensity at 535 nm with increasing site-marker concentration) (c) and (d) molecular docking picture shows the binding interaction of NBD-Bu with BSA.

To gain further insight into these phenomena, we have performed molecular docking studies and analyzed by PyMOL, which were consistent with competitive binding experiments with site markers. NBD-Bu structure, optimized using Gaussian 09, was further used for all docking studies. The probe binds to site-II with binding energy 6.64 kcal/mol, and the stabilization energy was found to be −7.32 kcal/mol.

After a detailed understanding of the specificity and binding phenomena of NBD-Bu with BSA, we investigated the validity and applicability of our findings using fluorescence microscopy. For this purpose, initially, the MTT assay was performed (Fig. S16) and IC_50_ value was found to be 46 *µ*M. Whereas, 70% of the cells were viable when incubated with 10 *µ*M dye concentration confirming the suitable working concentration for further cell imaging studies. Recent report^27^ suggests a key role of abnormal metabolism during cancer resulting in the disease itself, rather than uncontrolled cell proliferation. Therefore, different normal (CHO, and BHK-21) and cancer cell lines (B16F10), and HeLa) (Fig. 4) have been taken to distinguish the metabolic activity using NBD-Bu as a probe. The CHO and B16F10 cells were grown for 24h to reach 70% confluency followed by 6h of serum-starvation. Finally, they were incubated for 20 mins with 10 *µ*M NBD-Bu for imaging. The confocal microscopy images of CHO cells and B16F10 cells (Fig. 4a-b) show that the fluorescence intensity is considerably higher in the serum-starved cells (followed by 100 *µ*M BSA treatment) compared to the well-fed cells. This is evident from the intensity profiles obtained from the corresponding green channel images. The intensity in serum-starved CHO cells is ~4 times to that of untreated ones. On another hand, the intensity is only ~1.5 times in starved B16F10 cells compared to well-fed cells. This can be attributed to the higher uptake capacity of cancer cells due to higher metabolic activity compared to normal cells, which resulted in greater intensity even in the well-fed cells. This arguments towards the applicability of NBD-Bu as a metabolic marker to visualize the difference in metabolism in normal and cancer cells.

**Fig. 4.**
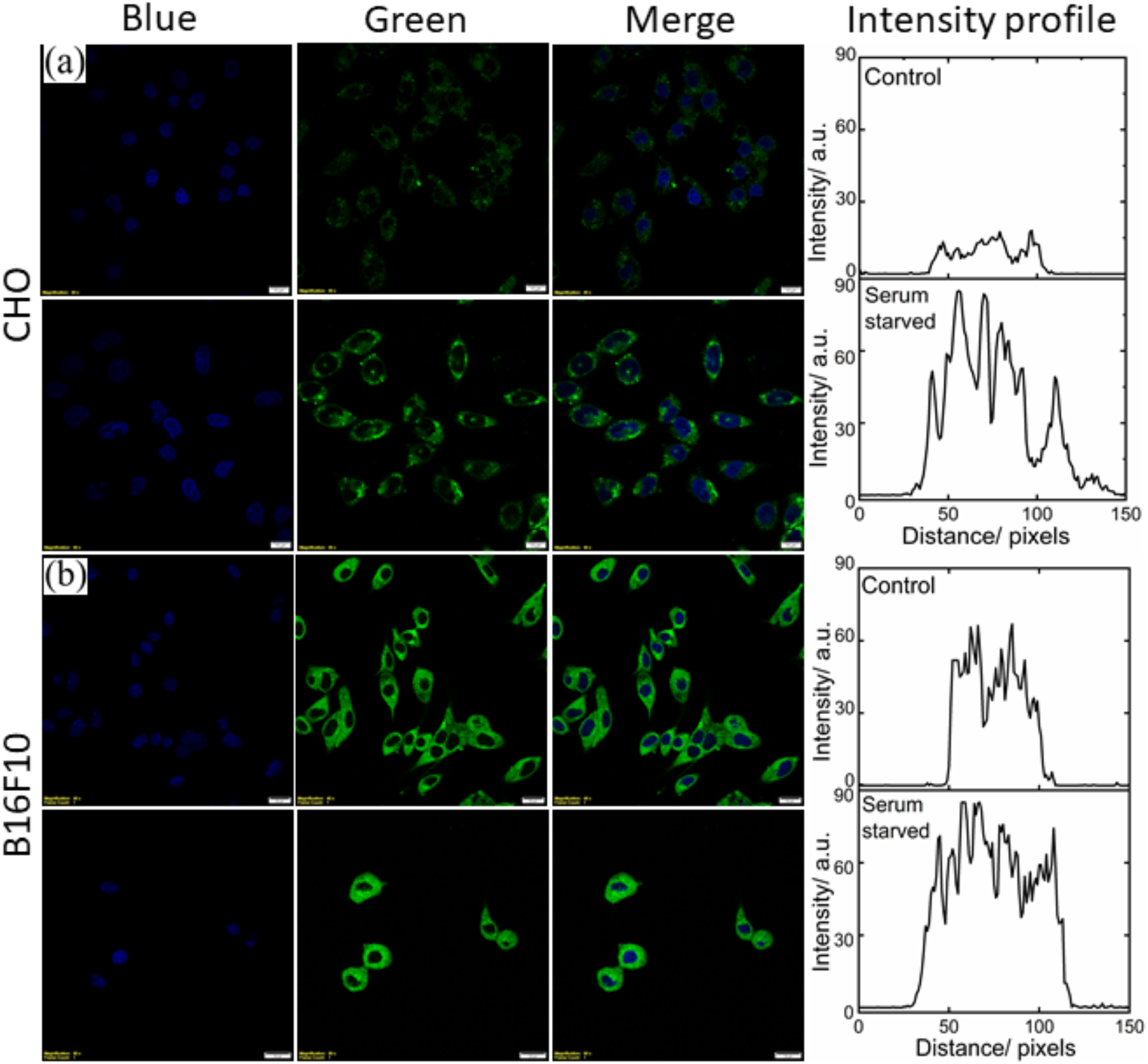
Live-cell confocal microscopy imaging of (a) B16F10 (scale bar 20 *µ*m) and (b) MCF-7 (scale bar 5 *µ*m) cell lines. The left blue panel is for nuclear staining using Hoechst 33342; middle panel for the green channel for NBD-Bu and right is merged of the blue and green channel. The corresponding average intensity obtained from the line profiles shows much higher fluorescence enhancement with serum-starved cells compared to untreated ones.

**Fig. 5.**
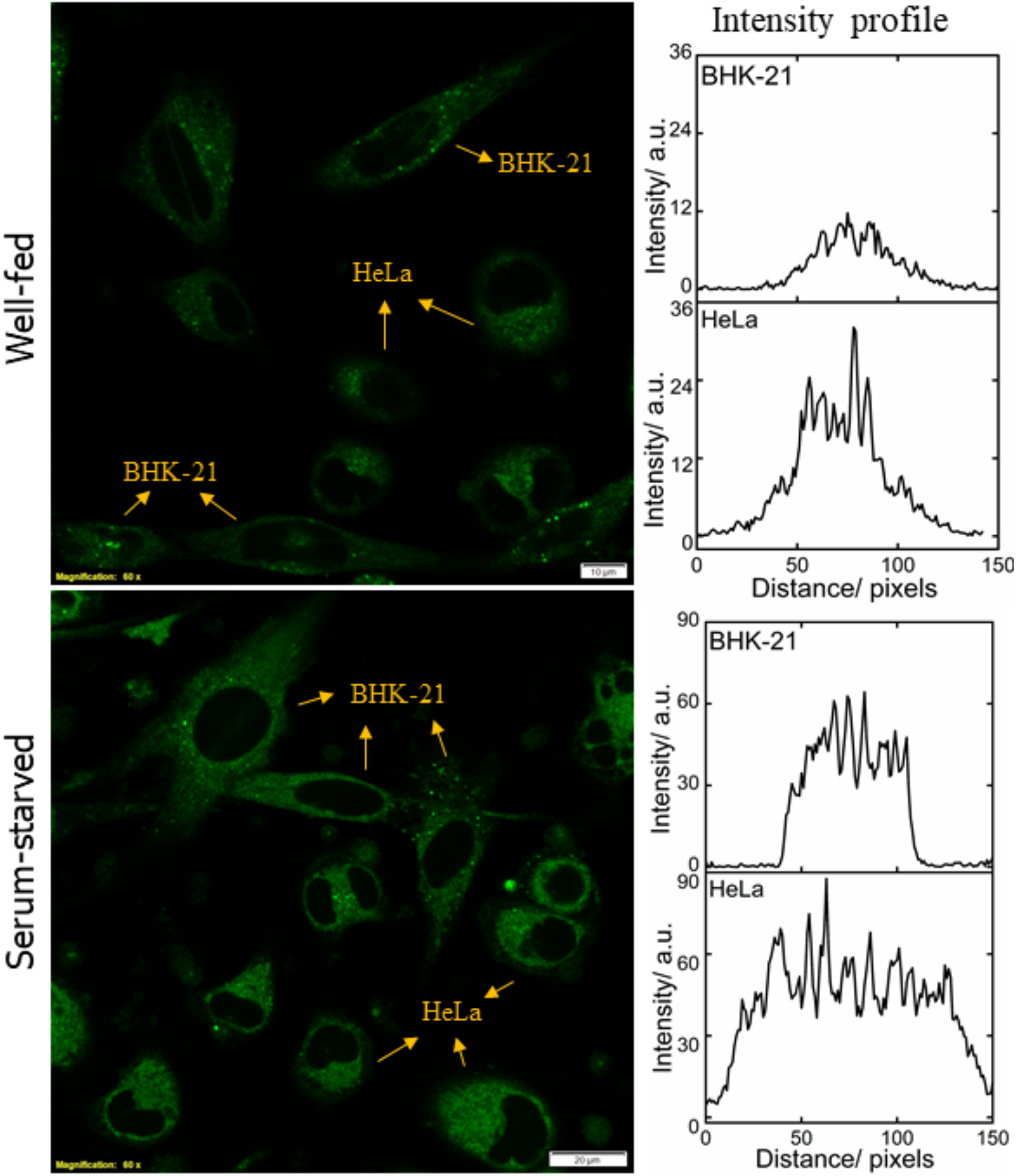
Live-cell confocal microscopy imaging of BHK-21 and HeLa cell lines in co-culture condition, (a) well-fed and (b) serum-starved cells and their corresponding intensity profile (averaged from 4 ROIs of each cell lines per image) shows the greater cellular uptake in cancer cells. Scale bar 20 *µ*M

The results obtained from the monoculture of normal and cancer cell lines were further verified with co-culture condition. BHK-21 and HeLa cells were seeded in 1:1 ratio and grown for 48h followed by 6h of serum-starvation. Serum-starved cells were additionally incubated with 100 *µ*M BSA for 30 min before further incubation for 20 min with 10 *µ*M NBD-Bu. Living up to our anticipation, the intensity recorded in well-fed HeLa cells was ~3.5 times compared to BHK-21 cells, whereas only ~1.5 times higher intensity was measured in serum-starved HeLa cells. These observations are well in synchronization with the monoculture results (*vide supra*).

## Conclusions

In summary, we have designed a NBD-based ICT probe for instantaneous and efficient detection of serum albumin in the presence of other biologically relevant proteins. The detailed spectroscopic studies revealed that the probe is binding to site-II of BSA and was further supported by the competitive binding and molecular docking studies. Finally, employing the binding phenomena, the difference in metabolic activity in non-cancerous and cancer cells have been well elucidated by the live-cell microscopic investigation in monoculture and co-culture conditions which showed that serum-starvation led to the enhancement of intensity in cancer cells compared to normal cells. We believe that this probe can be employed for the detection of abnormal serum albumin levels in the patient’s body fluids and as metabolic activity marker for diagnostics applications.

We would like to thank IISER Bhopal and Department of Biotechnology (DBT, BT/PR14498/NNT/28/880/2015), India for generous funding and infrastructural support. T.D. acknowledges IISERB and K.P. acknowledge the Council of Scientific & Industrial Research (CSIR), India for their doctoral fellowship.

