## supplemental file for "Cellular metabolic activity marker via selective turn-ON detection of serum albumin using an NBD-based fluoroprobe"

### EXPERIMENTAL SECTION

**Chemicals and Materials.** 4-Chloro-7-nitrobenzofurazan, warfarin, trypsin (from bovine pancreas), $\alpha$-Chymotrypsin (from bovine pancreas) and α-lactalbumin (from bovine milk) were purchased from Sigma-Aldrich (India). n-Butyl amine was purchased from Spectrochem (India). HSA, RNase A (from bovine pancreas, free from DNase), Catalase (from the bovine liver), $\alpha$-Amylase and DNase (from bovine pancreas) were purchased from HIMEDIA (USA). BSA was purchased from BIOMATIK, USA. All the solvents were of spectroscopic grade and were used without any further purification. Milli-Q grade water was used for the spectroscopic measurements with a resistivity of 18.2 MΩ·cm at 298 K.

**Synthesis and Characterization.** $^{1}H$ and $^{13}C\{^{1}H\}$ NMR spectra were recorded on Bruker 400 MHz spectrometers with operating frequencies of 100 MHz for $^{13}C$ using $CDCl_3$ as solvent and tetramethylsilane (TMS) as an internal standard. Chemical shifts ($\delta$) are reported in ppm relative to the residual solvent signal ($\delta$ = 7.26 for $^{1}H$ NMR and $\delta$ = 77.3 for $^{13}C$ NMR). HRMS data were recorded on MicrOTOF-Q-II mass spectrometer using Acetonitrile as a solvent.

To a solution of 4-Chloro-7-nitrobenzofurazan (100 mg, 0.501 mmol) in 10 mL methanol, butyl amine (99.03 $\mu$L, 1.002 mmol) was added dropwise over the course of 5 minutes. The reaction was performed in an inert atmosphere for 5 hours at room temperature. Then the solvent was removed under reduced pressure. The residue was purified by silica gel column chromatography using 30% - 40% mixture of EtOAc and Hexane as eluent and the yield was calculated to be 60%. $^{1}H$ NMR (400 MHz, $CDCl_3$) $\delta$: 8.5 (1H, d, $J$= 8.6 Hz), 6.18 (1H, d, $J$= 8.7 Hz), 3.5 (2H, q, $J$= 6.7 Hz), 1.8 (2H, m, $J$= 7.3 Hz), 1.51 (2H, dt, $J$= 14.8, 7.5 Hz), 1.02 (3H, t, $J$= 7.3 Hz). $^{13}C\{^{1}H\}$ NMR (400 MHz, $CDCl_3$) $\delta$: 144.26, 143.88, 136.50, 123.98, 98.50, 43.71, 30.54, 20.14, 13.68. HRMS (ESI) m/z $[M+H]^{+}$ calcd for $C_{10}H_{12}N_4O_3$ 237.0909 Da; found 237.0982 Da.

**Spectroscopic Measurements.** Steady-state absorption spectra were recorded on Cary 5000 UV-spectrophotometer. All steady-state fluorescence measurements were done on HORIBA Jobin Yvon Fluorolog spectrofluorimeter. All spectroscopic data were recorded by using a 1 cm path length quartz cuvette, $\lambda_{ex}$ =450 nm. Both excitation and emission slit were kept at 1 nm while recording the fluorescence spectra. Time-resolved fluorescence measurements were performed using a Hamamatsu MCP photomultiplier (R-3809U-50). The time-correlated single photon counting (TCSPC) setup consists of an Ortec 9327 pico-timing amplifier and using pulse Diode laser ($\lambda_{ex}$ =470 nm) with fwhm ~143 ps with a setup target 10,000 counts. The emission polarizer was positioned at the magic angle (54.7°) polarization w.r.t. the excitation polarizer. The single exponential fitting function was employed by iterative deconvolution method using supplied software DAS v6.2. The general form of the fitting function was

$$I(t)=I(0) \sum \alpha_i \exp(-t/\tau_i)$$

where I(t) and I(0) are the fluorescence intensity at time t and 0 respectively, t is the time $\alpha_i$ and $\tau_i$ are the contributing amplitude and its corresponding lifetime. The quality of the fitted data was judged from the reduced chi-squared value ($\chi^2$), calculated using the IBH software provided with the instrument. Circular dichroism spectra were measured using JASCO J-815 CD spectrometer with standard sensitivity (100 mdeg), 0.1 nm data pitch with scanning speed 200 nm $min^{-1}$.

**Optimization and Molecular Docking Studies.** The structure of NBD-Bu was optimized with Gaussian 09 with method RB3LYP/6-31G (Calculation type: FOPT)**.** The molecular docking studies have been done with SwissDock,[1] based on the EADock DSS engine, a web server for docking of small molecules on the target proteins. The PDB file of BSA (4F5S) and the optimized structure were used for the docking. The job was submitted to the server and obtained results were analyzed by PyMOL (v2.1.1) and UCSF Chimera[2] (v1.13) for visualization and understanding.

### Cell Culture and Imaging:

**Materials and Microscopy:** Dulbecco's Modified Eagle Medium (DMEM) and antibiotic cocktail were purchased from HiMedia (USA) and Fetal Bovine Serum (FBS) was purchased from Sigma Aldrich (USA). The cell imaging dishes were obtained from ibidi (Germany). All the confocal microscopy imaging were performed with an Olympus FV3000 Confocal Laser

Scanning Microscope (LSM). The image processing was done with the help of cellSens software (Olympus).

**Culture Method:** B16F10 and MCF 7 cells were obtained from NCCS, Pune, India and cultured in DMEM (phenol red free) containing 10% (v/v) FBS and 1% (v/v) antibiotic cocktail in 5% $CO_2$ at 37 ºC in incubator. For imaging purpose, cells were grown to 75% - 80% confluency in the 35 mm glass bottom imaging dishes (170 ± 5 $\mu$m). For co-culture experiment, BHK-21 and HeLa cells were seeded in 1:1 ratio and grown for 48h followed by 6h of serum-starvation. Serum-starved cells were additionally incubated with 100 $\mu$M BSA for 30 min before further incubation for 20 min with 10 $\mu$M NBD-Bu.

**Serum Starvation Experiment:** The growth medium was removed and washed twice with sterilized PBS buffer (pH 7.4, containing 5 mM $MgCl_2$). Then the cells were incubated with only DMEM (serum free) for 6 h. Thereafter, the cells were washed gently with PBS and further incubated with 100 $\mu$M BSA for 15 min at 37 ºC. After washing them again with PBS, all the cells, starved and non-starved ones, were incubated together with 8 $\mu$M Hoechst 33342 for nuclear staining and 10 $\mu$M NBD-Bu for 20 min at 37 ºC. Finally, they were washed and imaged under confocal LSM.

**Confocal Microscopy:** For fluorescence imaging, 405 nm (for Hoechst 33342 from ThermoFisher Scientific, USA) and 488 nm (for NBD-Bu) excitation lasers were used. For 405 nm and 488 nm excitation, the emission windows were kept at 430-470 nm and 500-600 nm, respectively. The confocal aperture was kept at 0.81 Airy Disk (AU) while the dwell time is 10 $\mu$s/pixel. The laser power, gain and offset were kept same for all.

**Image Analysis:** The post-processing was performed using ImageJ (Fiji) software[3] (v 1.52n). The line profiles of the selected ROIs were plotted as grayscale intensity vs. distance. These values were averaged to finally obtain an intensity profile.

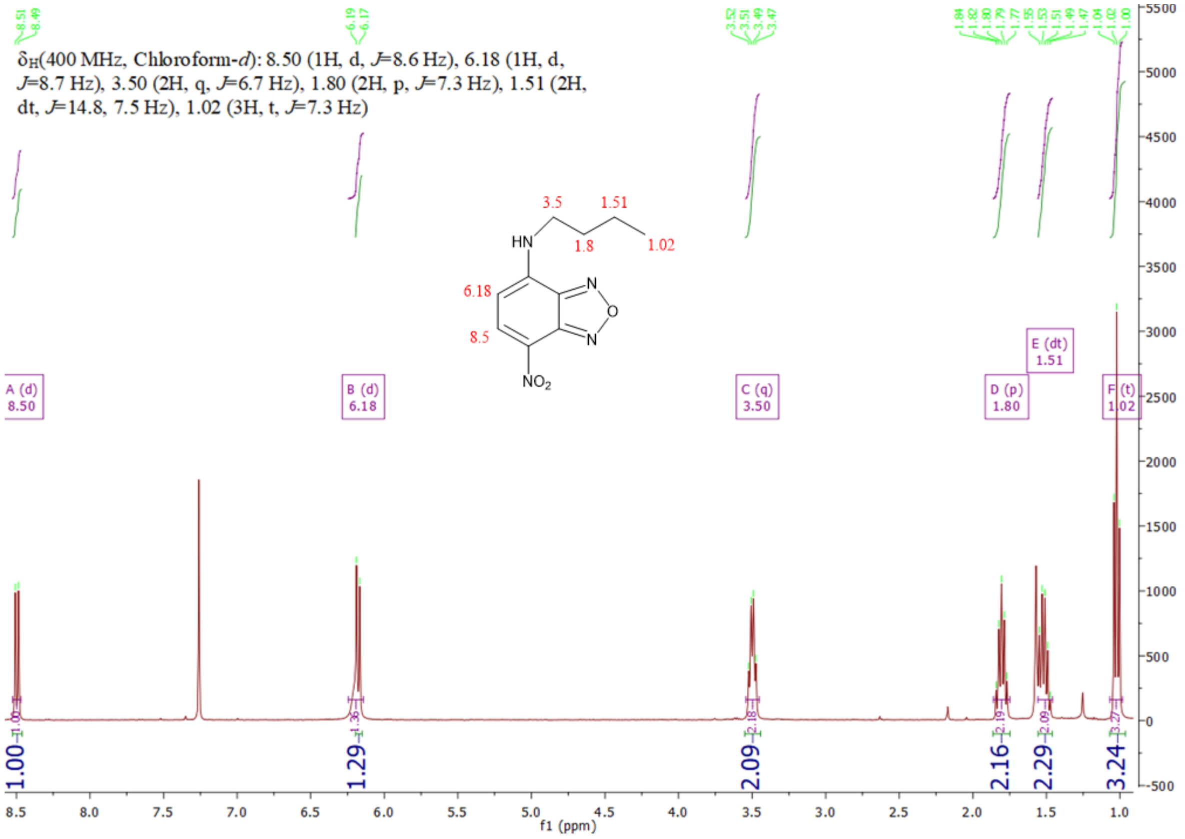


**Fig. S1** $^{1}$H NMR spectra (400 MHz) of NBD-Bu in $CDCl_3$

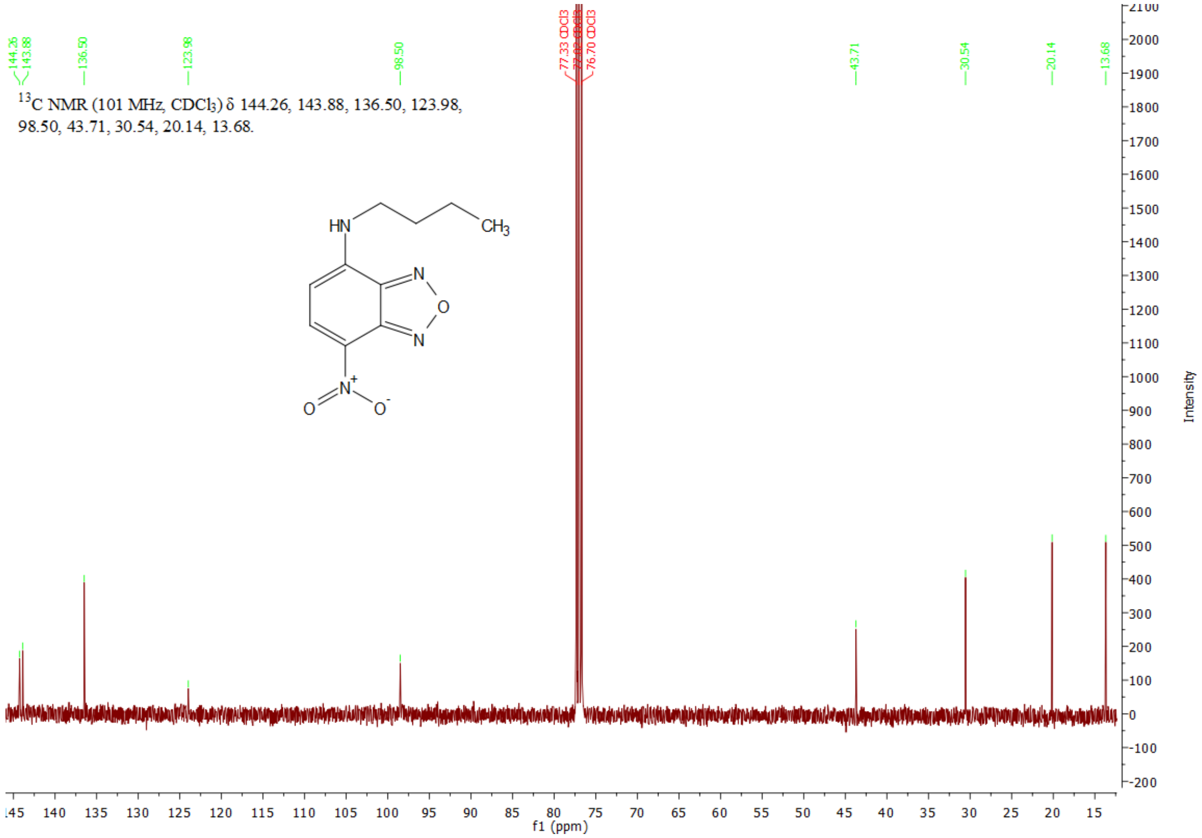


**Fig. S2** $^{13}C\{^{1}H\}$ NMR spectra (126 MHz) of NBD-Bu in $CDCl_3$

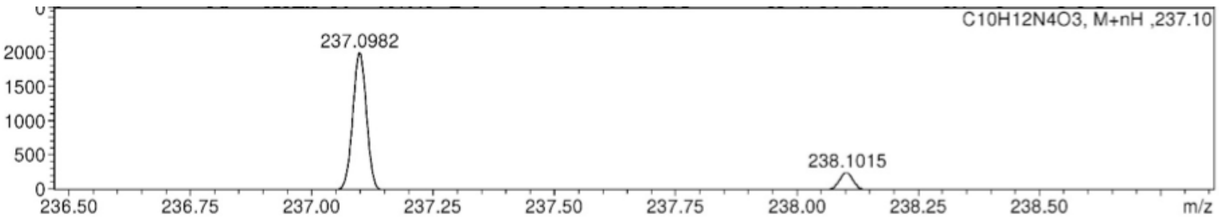


**Fig. S3** High-Resolution Mass Spectrum of NBD-Bu

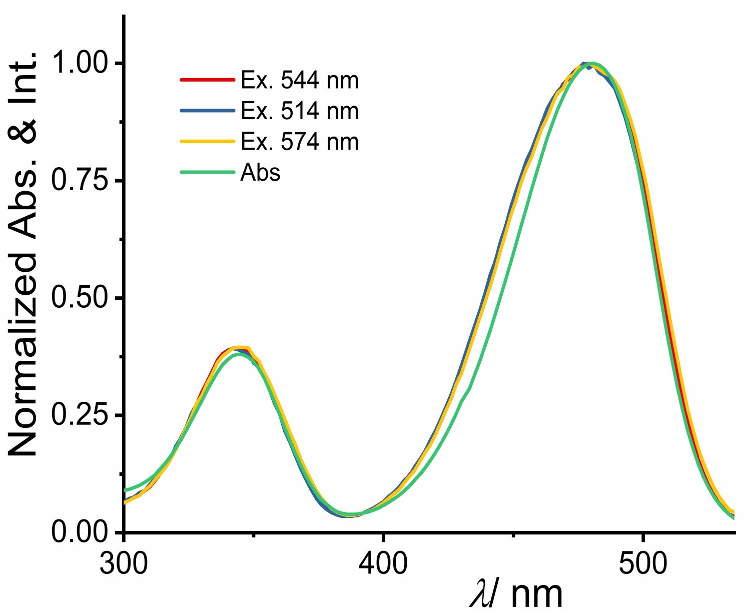


**Fig. S4** Testing the optical purity of NBD-Bu using UV-Vis. and excitation spectra in water (~pH 7.0).

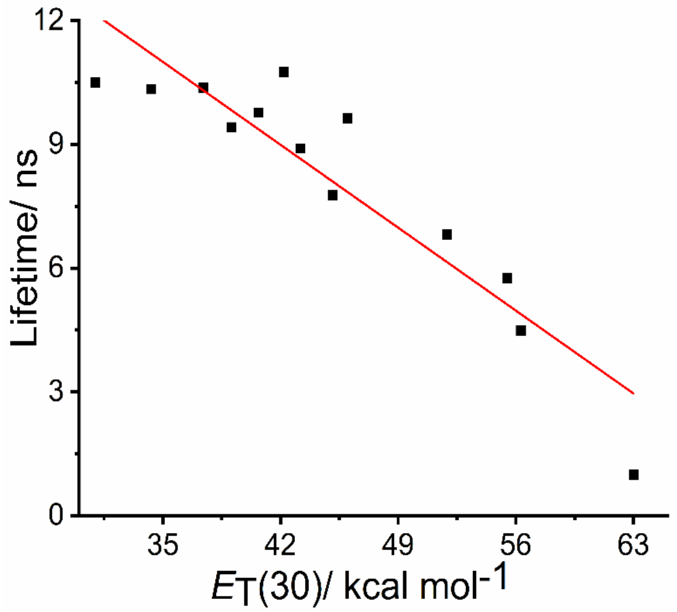


**Fig. S5** The linear variation of the lifetime of NBD-Bu (10 $\mu$M) with solvent polarity parameter, $R^2 = 0.85$

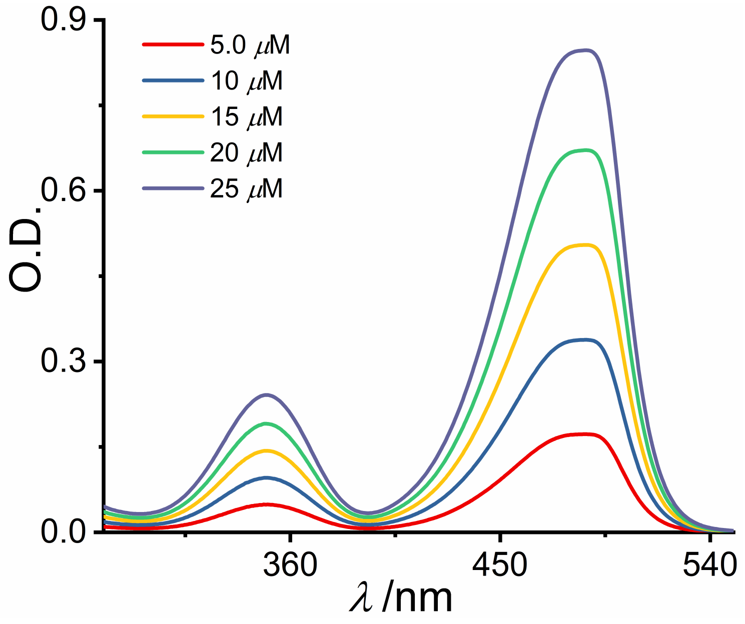


**Fig. S6** Absorption spectra of NBD-Bu in an aqueous medium in a different concentration

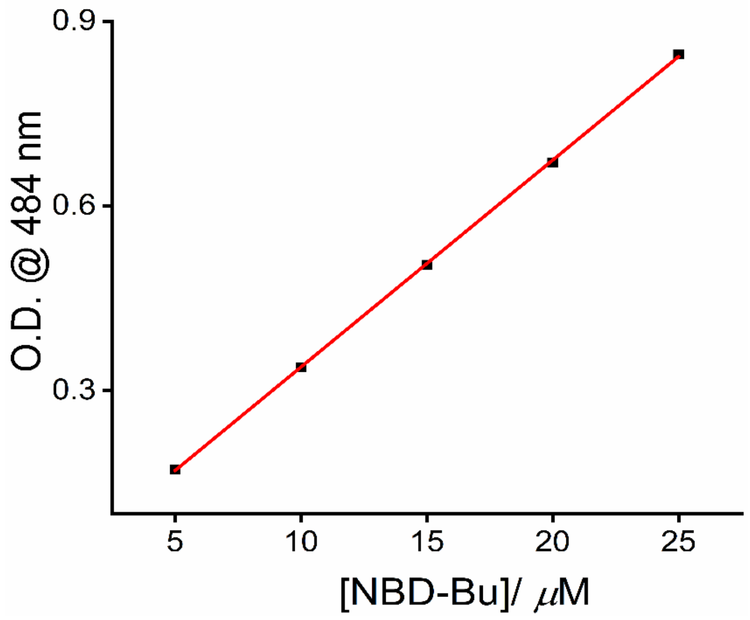


**Fig. S7** The plot of O.D. against increasing concentration of NBD-Bu at 484 nm ($R^2$= 0.99)

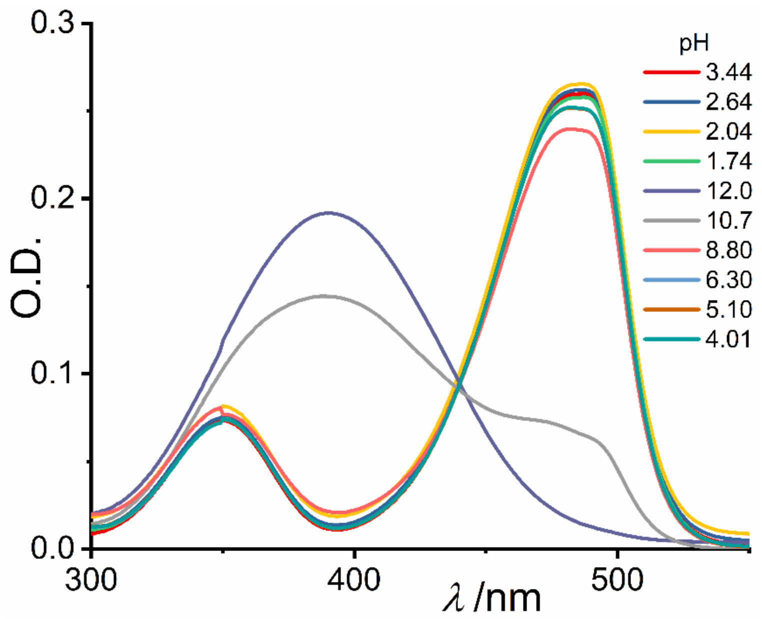


**Fig. S8** pH titration of NBD-Bu (10 $\mu$M)

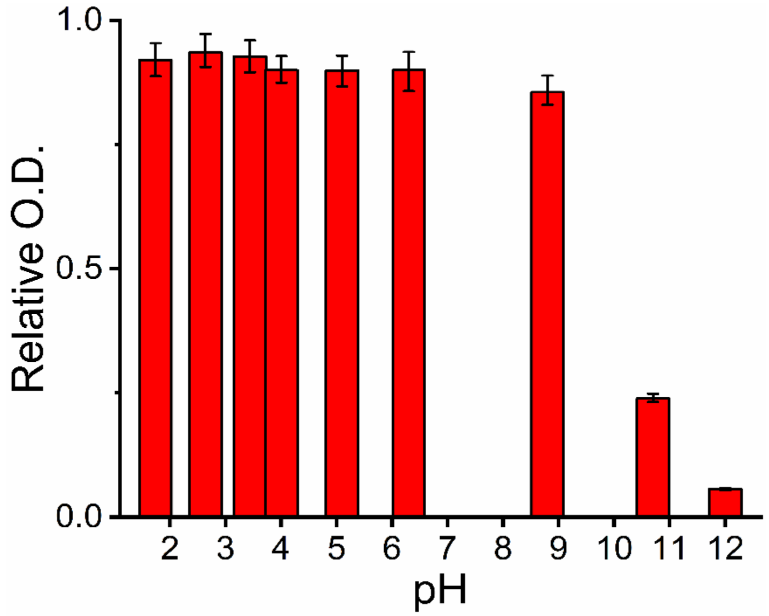


**Fig. S9** The absorbance of NBD-Bu (10 $\mu$M) in different pH

**Table S1:** Solvent-dependent photophysical properties of NBD-Bu

| Solvent | $\lambda_{abs}$/nm | $\lambda_{em}$/nm | Average lifetime/ ns | Quantum Yield[a] | $E_T(30)$[4] |
|---|---|---|---|---|---|
| Acetone | 462 | 528 | 10.75 | 0.85 | 42.2 |
| ACN | 465 | 533 | 9.63 | 0.82 | 46 |
| Benzene | 453 | 520 | 10.34 | 1.135 | 34.3 |
| Chloroform | 452 | 527 | 9.41 | 1.05 | 39.1 |
| DCM | 456 | 523 | 9.77 | 0.65 | 40.7 |
| DMSO | 481 | 544 | 7.77 | 0.66 | 45.1 |
| DMF | 473 | 538 | 8.90 | 0.86 | 43.2 |
| Ethanol | 466 | 536 | 6.81 | 0.62 | 51.9 |
| Ethylene glycol | 476 | 552 | 4.48 | 0.30 | 56.3 |
| Hexane | 448 | 520 | 10.50 | 0.64 | 31 |
| Methanol | 467 | 539 | 5.76 | 0.35 | 55.5 |
| THF | 461 | 524 | 10.37 | 0.87 | 37.4 |
| Water (PBS) | 484 | 565 | 0.99 | 0.06 | 63 |
| BSA (100 $\mu$M, PBS) | 484 | 535 | 6.1 | 0.153 | ~42 (polarity is like acetone) |

[a]Quantum yields were measured w.r.t. fluorescein dye in 0.1(N) NaOH.[3]

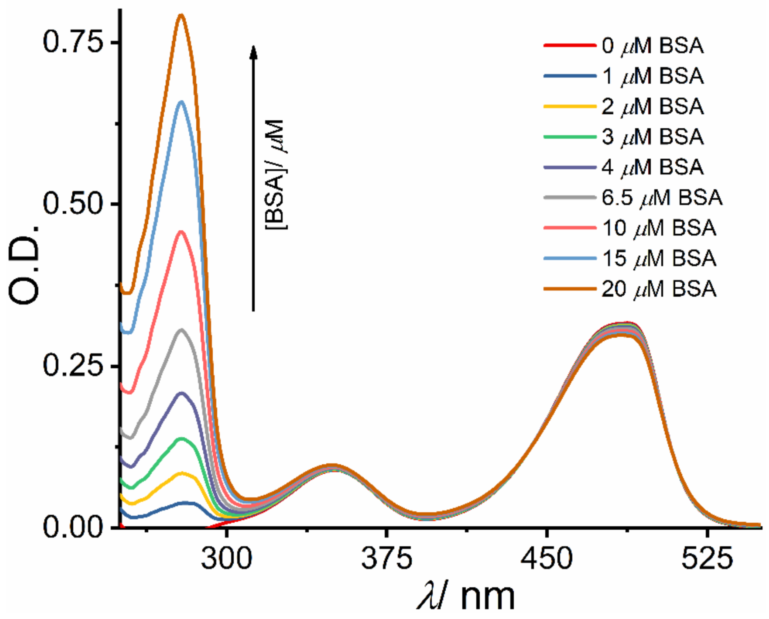


**Fig. S10** UV-Vis. titration of NBD-Bu (10 $\mu$M) with increasing BSA concentration in PBS

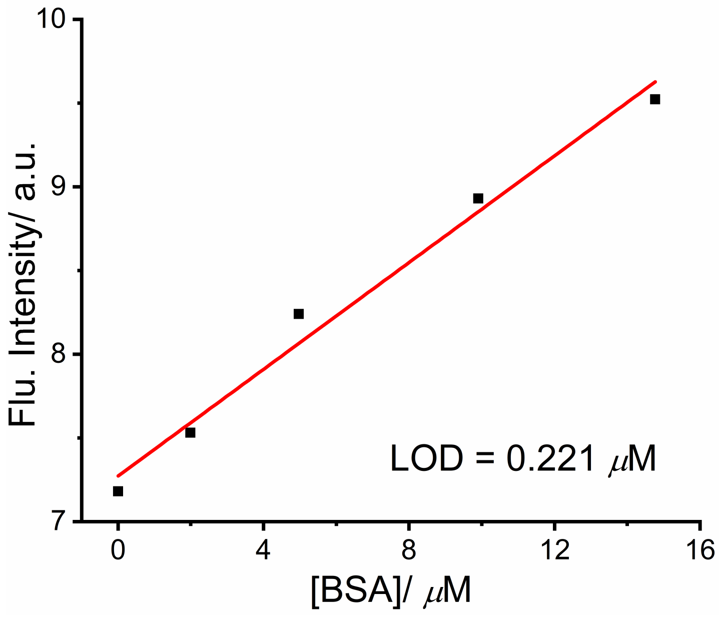


**Fig. S11** Estimation of the limit of detection (3$\sigma$/slope) from the calibration curve with BSA concentration ranging from 0-15 $\mu$M, $R^2$= 0.98

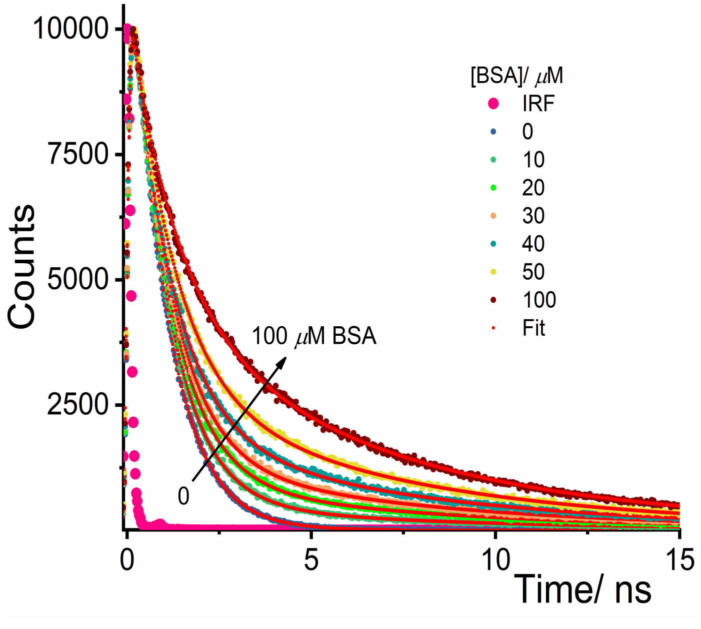


**Fig. S12** Increase in the fluorescence lifetime of NBD-Bu with increasing BSA concentration in PBS

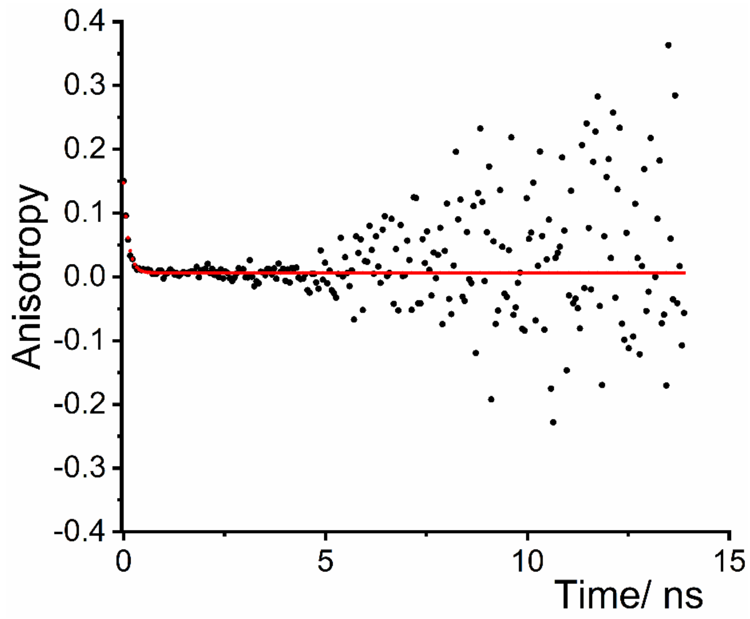


**Fig. S13** Fluorescence anisotropy of 10 $\mu$M NBD-Bu in PBS (pH 7.4)

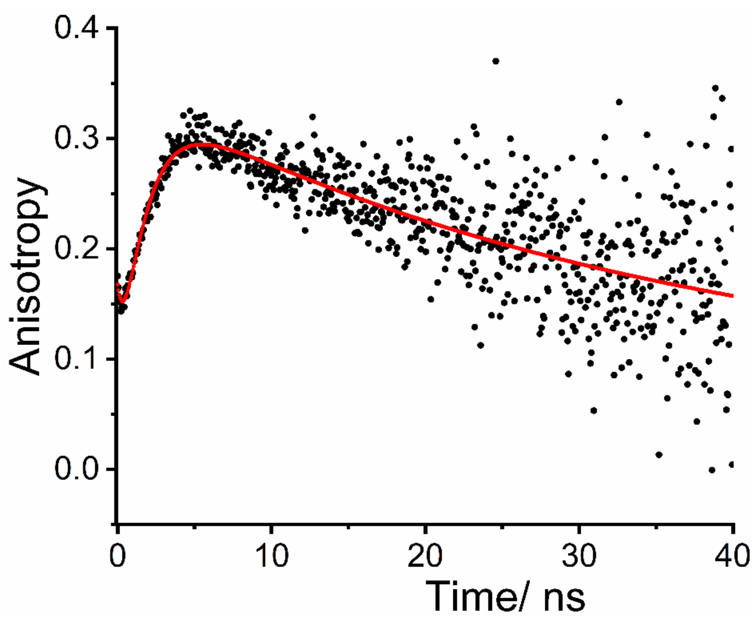


**Fig. S14** Fluorescence anisotropy of 10 $\mu$M NBD-Bu in presence of 100 $\mu$M BSA in PBS (pH 7.4)

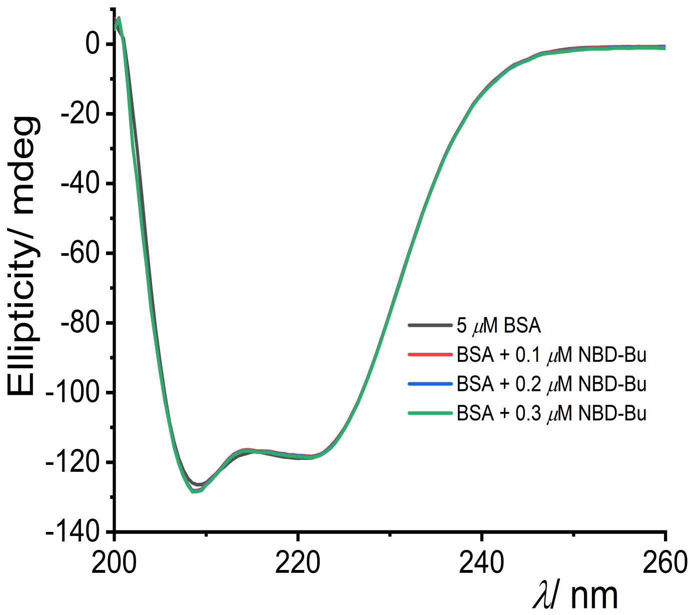


**Fig. S15** Circular dichroism spectra of BSA (5 $\mu$M) with increasing concentration of NBD-Bu in PBS

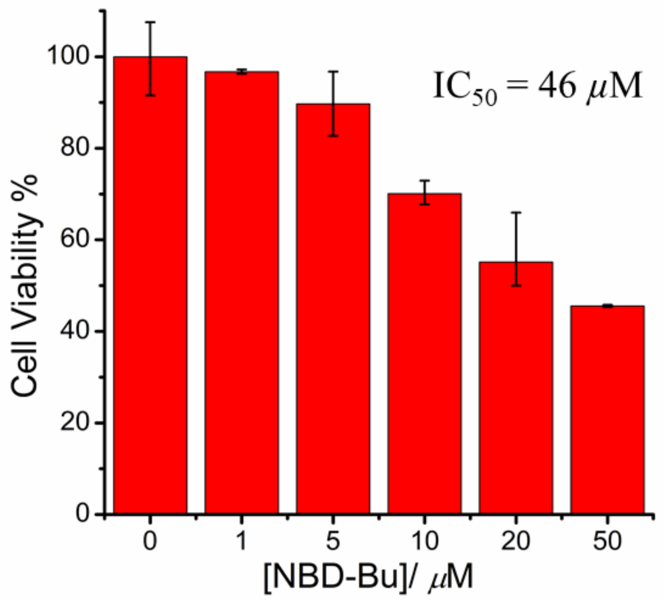


**Fig. S16** MTT assay to assess the cell viability of B16F10 cells against the different concentration of NBD-Bu
